# IL-1β–mediated inflammatory signaling drives ineffective erythropoiesis in early-stage myelodysplastic syndromes

**DOI:** 10.1101/2023.09.28.560018

**Authors:** Vera Adema, Irene Ganan-Gomez, Feiyang Ma, Juan Jose Rodriguez-Sevilla, Kelly Chien, Hui Yang, Natthakan Thongon, Rashmi Kanagal-Shamanna, Sanam Loghavi, Guillermo Montalban-Bravo, Danielle Hammond, Yiqian Gu, Roselyn Tan, Lin Tan, Philip Lorenzi, Gheath Al-Atrash, Karen Clise-Dwyer, Rafael Bejar, Matteo Pellegrini, Guillermo Garcia-Manero, Simona Colla

## Abstract

Myelodysplastic syndromes (MDS) are a group of incurable hematopoietic stem cell (HSC) neoplasms characterized by peripheral blood cytopenias and a high risk of progression to acute myeloid leukemia. MDS represent the final stage in a continuum of HSCs’ genetic and functional alterations and are preceded by a premalignant phase, clonal cytopenia of undetermined significance (CCUS). Dissecting the mechanisms of CCUS maintenance may uncover therapeutic targets to delay or prevent malignant transformation.

Here, we demonstrate that *DNMT3A* and *TET2* mutations, the most frequent mutations in CCUS, induce aberrant HSCs’ differentiation towards the myeloid lineage at the expense of erythropoiesis by upregulating IL-1β–mediated inflammatory signaling and that canakinumab rescues red blood cell transfusion dependence in early-stage MDS patients with driver mutations in *DNMT3A* and *TET2*.

This study illuminates the biological landscape of CCUS and offers an unprecedented opportunity for MDS intervention during its initial phase, when expected survival is prolonged.

## MAIN

Early therapeutic interventions after cancer detection can extend patient survival^1^. A key challenge is to dissect the biological mechanisms of preneoplastic disease states that precede tumorigenesis to slow down the evolution of the disease and/or overcome disease-related comorbidities^1^.

Here, to uncover early therapeutic approaches that may improve the clinical course of myelodysplastic syndromes (MDS), we sought to unravel the molecular and cellular mechanisms underlying clonal cytopenias of undermined significance (CCUS), a precursor state of MDS characterized by reduced clonal burden, lower genetic complexity, and milder peripheral blood cytopenias^2^.

Recognizing that CCUS is an aging-related disorder, we first dissected the way in which CCUS-related mutations affect the function of aging hematopoietic stem and progenitor cell (HSPC) compartment. We performed single-cell RNA-sequencing (scRNA-seq) analysis of Lin^-^ CD34^+^ HSPCs isolated from the BM of young healthy donors (yHDs), elderly HDs (eHDs), and patients with CCUS harboring *DNMT3A* and/or *TET2* mutations (Supplemental Table 1), which are the most commonly mutated genes in CCUS^3^. We identified 16 cellular clusters (Fig. 1a) driven by different cellular differentiation profiles, which we defined based on the differential expression of previously validated lineage-specific transcription factors (TFs) and cellular markers among the clusters^4–7^ (Supplemental Fig. 1a and Supplemental Table 2).

**Fig. 1.**
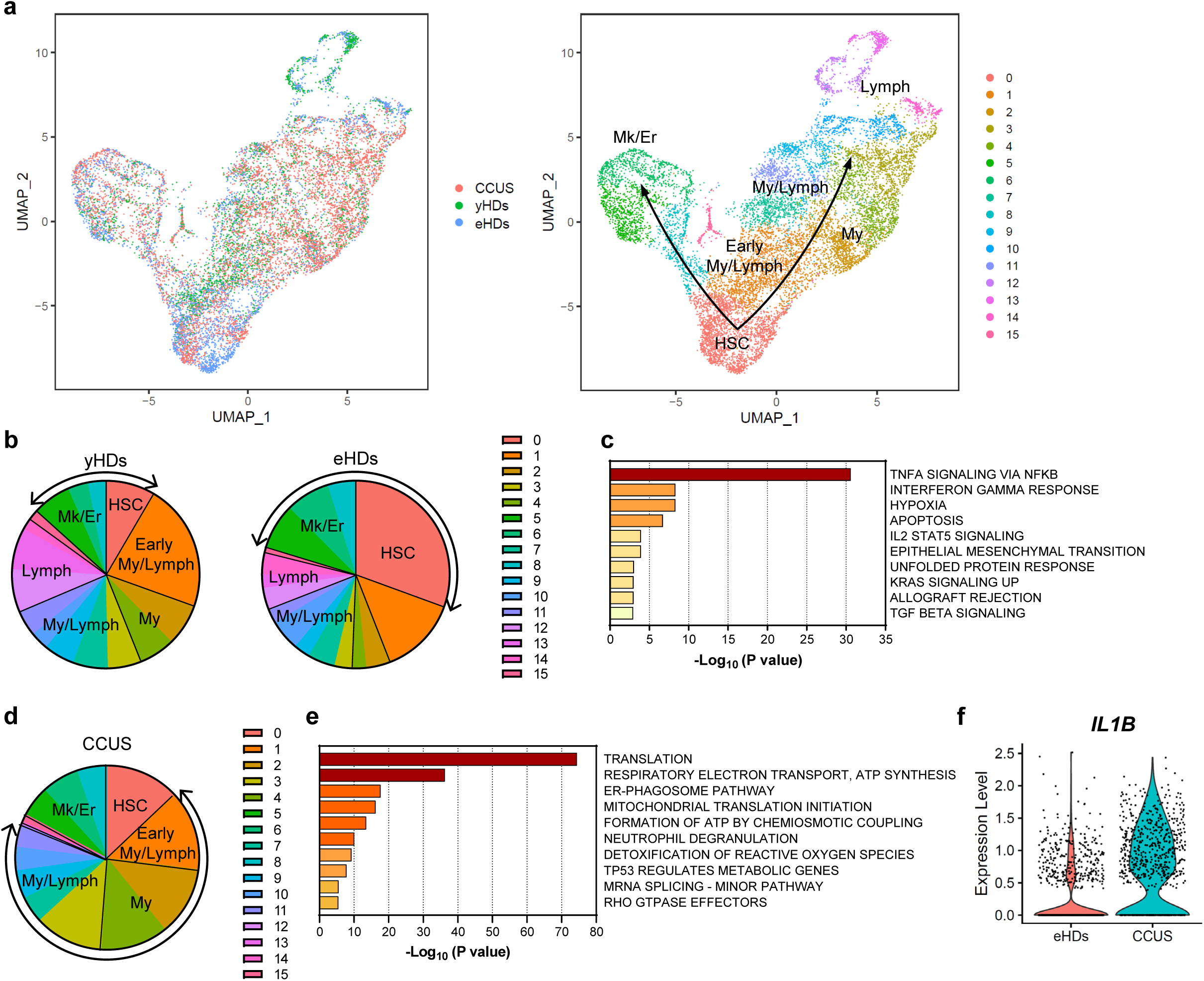
*DNMT3A* and *TET2* mutations transcriptionally rewire the HSCs’ aging phenotype. **(a)** UMAP plot of scRNA-seq data from Lin^-^CD34^+^ cells isolated from 3 yHDs (n = 2,823), 4 eHDs (n = 3,655), and 3 CCUS patients (n = 6,101). Each dot represents one cell. Different colors represent the sample (left) and cluster (right) identities. Lymph, lymphoid; Mk, megakaryocytic; Er, erythroid; My, myeloid; HSC, hematopoietic stem cell. **(b)** Cluster distribution of the Lin^-^CD34^+^ cells isolated from yHDs and eHDs shown in Fig. 1a, represented as the percentage of cells in each cluster. Clusters are grouped based on the cells’ lineage annotation. **(c)** Pathway enrichment analysis of genes that were significantly (*P* adj ≤ 0.05) upregulated in HSCs from eHDs compared with those in HSCs from yHDs. The top 10 Hallmark gene sets are shown. **(d)** Cluster distribution of Lin^-^CD34^+^ cells from CCUS patients, represented as the percentage of cells in each cluster. Clusters are grouped based on the cells’ lineage annotation. **(e)** Pathway enrichment analysis of genes that were significantly (*P* adj ≤ 0.05) upregulated in HSCs from CCUS patients compared with those in HSCs from eHDs. The top 10 Reactome gene sets are shown. **(f)** Expression of *IL-1β* in HSCs from eHDs and CCUS patients. Each dot represents the expression level of a single cell (*P* adj = 1.70 × 10^-71^).

To dissect the changes in lineage specification that occur during physiological aging, we compared the distribution of yHD and eHD HSPCs in the distinct hematopoietic clusters. We found that, in contrast to the long-standing observation that aging HSPCs are mostly myeloid-biased^8,9^, HSPCs from eHDs, compared with those from yHDs, had increased HSC frequencies (characterized by the highest expression of *MLLT3*, *HLF*, *MEG3,* and *CDKN1C*; Supplemental Fig. 1b) and megakaryocytic-erythroid transcriptional signatures but reduced early and late myeloid/lymphoid signatures (Fig. 1b). These results were confirmed by immunophenotypic analyses of Lin^-^CD34^+^ cells using previously defined HSPC surface markers^10^ (Supplemental Fig. 1c). Further differential transcriptomic analyses across each cluster showed that HSCs from eHDs were characterized by a significant upregulation of genes involved in inflammation-mediated signaling, including the tumor necrosis factor alpha (TNF-α)-induced activation of nuclear factor kappa-light-chain-enhancer of activated B cells (NF-κB) and the cellular response to interferon gamma (IFN-γ; Fig. 1c, Supplemental Fig. 1d,e, and Supplemental Table 3). Importantly, the anti-apoptotic and NF-κB’s downstream effector gene *MCL1*^11,12^ was also significantly upregulated in HSCs from eHDs (*P* adj = 3.71 ×10^-14^; Supplemental Fig. 1f), which aligns with previous findings that the TNF-α–mediated activation of NF-κB is a major survival pathway in aged HSCs^13^. Similar results were obtained from scRNA-seq analysis of highly purified Lin^-^CD34^+^CD38^-^CD90^+^CD45RA^-^ HSCs isolated from yHDs and eHDs (Supplemental Fig. 1g-j).

Next, we sought to dissect the biological effects of CCUS-related mutations on the aging HSPC compartment. Differential analysis of cell composition revealed that, compared with those from eHDs, HSPCs from patients with CCUS had a predominantly myeloid-biased differentiation trajectory, which was characterized by an increased frequency of early myeloid/lymphoid progenitors and committed myeloid progenitors (Fig. 1d). However, flow cytometry of samples from a large cohort of eHDs and CCUS patients showed that CCUS patients have significantly fewer numbers of CD34^+^CD38^-^ HSCs and CD34^+^CD38^+^ hematopoietic progenitor cells in the BM (Supplemental Fig. 1k), which suggests that myeloid priming in CCUS HSPCs is the result of aberrant differentiation of HSCs towards the myeloid lineage rather than a numerical expansion of downstream myeloid progenitor cells. Consistent with this observation, we observed that genes involved in translation, respiratory electron transport, and mitochondrial translation initiation were significantly upregulated in HSCs from CCUS patients compared with those in HSCs from eHDs (Fig. 1e and Supplemental Fig. 1l-n), which underscores the metabolic activation of these cells in CCUS. Targeted ion chromatography–mass spectrometry analysis of CD34^+^ HSPCs isolated from the BM of CCUS patients and eHDs confirmed that CCUS CD34^+^ HSPCs had significant a upregulation of intermediates involved in the tricarboxylic acid cycle pathway and downregulation of those involved in glycolysis (Supplemental Fig. 1o). These results suggest that HSCs, which rely on anaerobic glycolysis to maintain their quiescent state in homeostatic conditions^14^, switch to mitochondrial oxidative metabolism in CCUS to meet the high-energy demands necessary to sustain activated myeloid differentiation^15,16^. Further differential transcriptomic analysis of eHD and CCUS HSCs showed that the expression of the pro-inflammatory cytokine interleukin 1 beta (*IL-1*β) was significantly upregulated in CCUS HSCs (*P* adj = 1.70 × 10^-71^; Fig. 1f and Supplemental Table 4). Notably, HSCs also expressed the IL-1β receptor (*IL-1R1*) (Supplemental Fig. 1p). In light of previous findings in mice that chronic IL-1β exposure drives HSC differentiation towards myelopoiesis at the expense of erythropoiesis and lymphopoiesis^17^, these results may explain why CCUS HSCs are myeloid-primed and metabolically activated. Interestingly, the expression levels of other cytokines, such as *CXCL2*, *CXCL3,* and *MIF*, which play a role in the pathogenesis of cardiovascular disease^18^, were also significantly increased in CCUS HSCs (Supplemental Fig. 1q).

An inflammatory environment enhances clonal dominance mutant HSPCs over their normal HSPC counterparts during early-stage clonal hematopoiesis^19^. To evaluate whether HSPCs’ extrinsic inflammatory mechanisms contribute to mutant HSPCs’ maintenance in CCUS, we performed scRNA-seq analysis of BM mononuclear cells (MNCs) isolated from yHDs and eHDs and from CCUS patients with *DNMT3A* and/or *TET2* mutations. Our analysis identified 26 cellular clusters including all major BM cell types, which we defined according to known cellular surface markers and lineage potential^20^ (Fig. 2a, Supplemental Fig. 2a, and Supplemental Table 5). Differential analysis of BM cell type compositions revealed higher frequencies of HSPCs and myelomonocytic cells in BM from eHDs than in BM from CCUS patients (Supplemental Fig. 2b), which aligns with our analyses of Lin^-^CD34^+^ cells and the cytopenias observed in the peripheral blood of these CCUS patients (Supplemental Table 1). Compared with those from yHDs, myelomonocytic cells from eHDs overexpressed genes involved in NF-κB signaling and inflammatory response pathways, including *IL-1β* (Supplemental Fig. 2c,d and Supplemental Table 6), which is consistent with the observation that aged myelomonocytes have increased inflammatory cytokine production^21^. CCUS myelomonocytic cells, which were highly metabolically active, exacerbated inflammatory signaling and overexpressed genes involved in IFN and complement responses (Fig. 2b and Supplemental Table 7). Importantly, the expression of *CASP1*, which cleaves pro–IL-1β to an active secreted cytokine^22,23^ (Supplemental Fig. 2e), and that of genes belonging to the S100A family of pro-inflammatory factors, which induce NF-κB pathway activation and promote the production of pro-inflammatory mediators^24,25^, including IL-1β, were also significantly upregulated in myelomonocytic cells from CCUS patients compared with those in myelomonocytic cells from eHDs (Supplemental Fig. 2f).

**Fig. 2.**
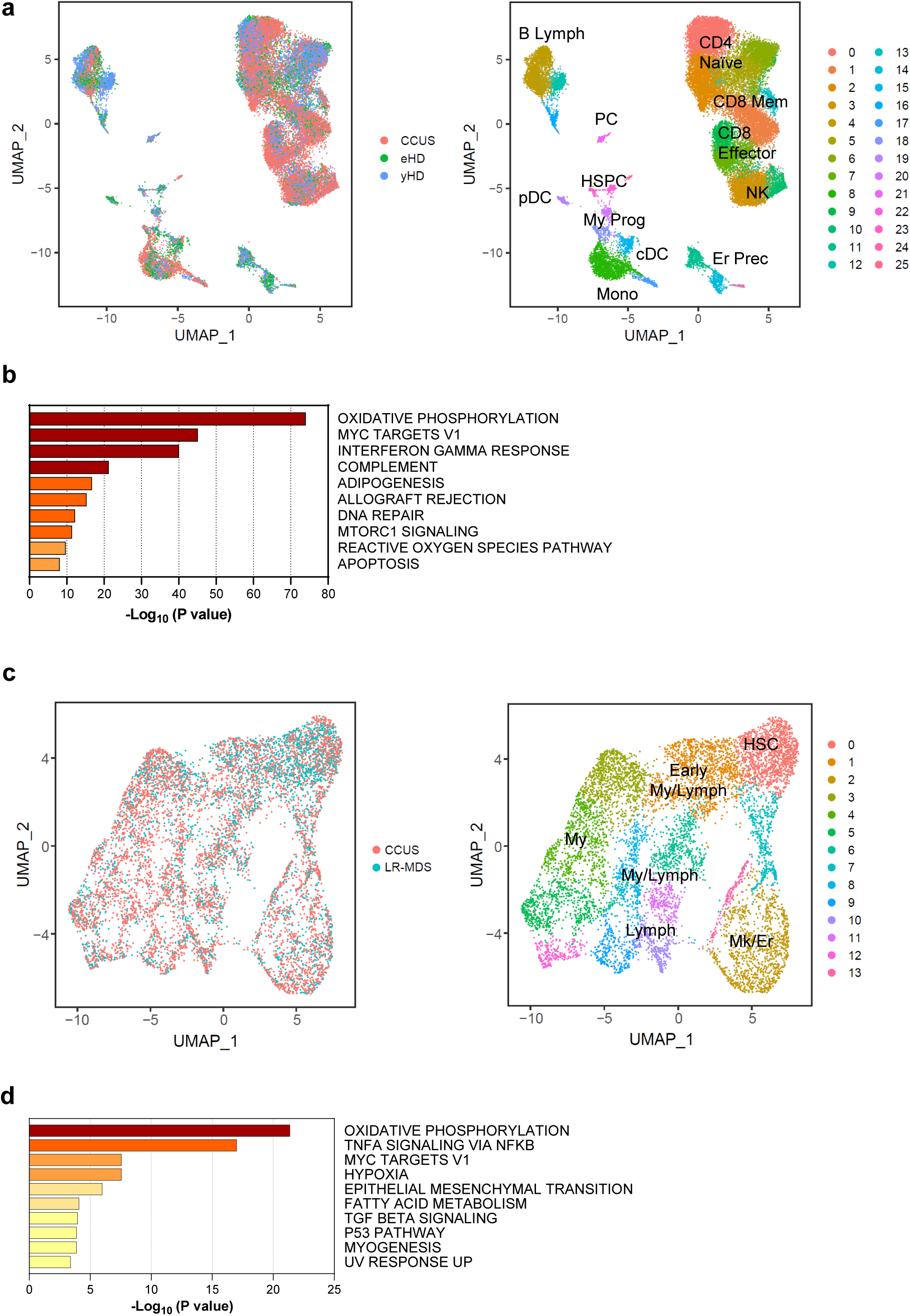
IL-1β–mediated inflammatory signaling is an initiation event in the pathogenesis of CCUS. **(a)** UMAP plot of scRNA-seq data from BM MNCs isolated from 2 yHDs (n = 9,375), 2 eHDs (n = 5,605), and 4 CCUS patients (n = 16,942). Each dot represents one cell. Different colors indicate the sample origin (left) and transcriptional cluster identity (right) for each cell. Clusters are annotated based on their lineage commitment. Lymph, lymphoid; Mem, memory; PC, plasma cell; HSPC, hematopoietic stem and progenitor cell; pDC, plasmacytoid dendritic cell; NK, natural killer; My, myeloid; Prog, progenitor; cDC, classic dendritic cell; Er, erythroid; Prec, precursor; Mono, monocytic. **(b)** Pathway enrichment analysis of genes that were significantly (*P* adj ≤ 1 × 10^-5^) upregulated in myelomonocytic cells from CCUS patients compared with those in myelomonocytic cells from eHDs. The top 10 Hallmark gene sets are shown. **(c)** UMAP plot of scRNA-seq data from Lin^-^CD34^+^ cells isolated from the BM of 3 CCUS patients (n = 5,822) and 3 LR-MDS patients (n = 2,970). Each dot represents one cell. Different colors represent the sample (left) and cluster (right) identities. HSC, hematopoietic stem cell; My, myeloid; Lymph, lymphoid; Mk, megakaryocytic; Er, erythroid. (**d**) Pathway enrichment analysis of genes that were significantly (*P* adj ≤ 0.05) upregulated in HSCs from LR-MDS patients compared with those in HSCs from CCUS patients. The top 10 Hallmark gene sets are shown.

To evaluate whether IL-1β–mediated inflammation increased in patients with lower-risk MDS (LR-MDS) and potently contributed to disease progression, we compared the expression profile of Lin^-^CD34^+^ cells isolated from patients with CCUS with *DNMT3A* and/or *TET2* mutations with that of Lin^-^CD34^+^ cells isolated from LR-MDS patients whose disease had mutations in the same genes along with multiple other molecular alterations (Fig. 2c and Supplemental Tables 1,8).

Differential expression analysis of HSCs showed that disease progression was associated with an exacerbation of CCUS transcriptomic alterations, including further activation of oxidative phosphorylation and NF-κB–mediated inflammatory signaling pathways (Fig. 2d). Compared with CCUS HSCs, LR-MDS HSCs overexpressed inflammatory cytokine genes involved in the activation of the CXCR2 axis, including *MIF*, *CXCL8* (*IL8*), and *CXCL2*, but not those involved in the IL-1β signaling pathway (Supplemental Fig. 2g and Supplemental Table 9). These data are consistent with previous findings that the CXCR2 axis plays a role in the pathogenesis of MDS^26^ and cardiovascular disease^27^, the risk of which is high in LR-MDS patients^28^. Surprisingly, differential single-cell expression analysis of monocytes from patients with CCUS and BM MNCs from LR-MDS patients (Supplemental Fig. 2h) showed that the expression of genes involved in the IL-1β–mediated inflammatory pathway, including *IL-1*β and *CASP1* (Supplemental Fig. 2i and Supplemental Table 10), decreased significantly as the disease progressed to LR-MDS and acquired clonal diversity induced by mutations other than those affecting *DNM3TA* or *TET2*. These results suggest that IL-1β–mediated inflammatory signaling is an initiation event in the pathogenesis of clonal hematopoiesis but does not drive the progression of CCUS to MDS.

Based on these findings, we hypothesized that targeting IL-1β signaling with canakinumab, a monoclonal antibody that inhibits the binding of IL-1β to IL-1R1^29^, overcomes aberrant HSPC differentiation and inhibits the production of other inflammatory cytokines from the BM microenvironment only in patients with lower genetic complexity, such as single *DNMT3A* and/or *TET2* mutations. To test this hypothesis, we analyzed the responses of LR-MDS patients (n = 25; Supplemental Table 11) with symptomatic anemia and/or transfusion dependence who were enrolled in a phase II open label clinical trial of canakinumab and for whom at least one line of prior therapy had failed (NCT04239157). The only 2 patients who achieved stable erythroid hematological improvement (lasting > 10 months) and became red blood cell transfusion–independent (patients UPN-01 and UPN-02) each had a single driver somatic mutation in *TET2* or *DNMT3A,* as the majority of patients with CCUS does (Supplemental Fig. 3a). As a proof-of-concept, scRNA-seq analysis of Lin^-^CD34^+^ HSPCs from a patient with a *DNMT3A* mutation (UPN-02), who had stable erythroid hematological improvement after 2 cycles of canakinumab (Fig. 3a), revealed that canakinumab increased HSCs’ differentiation towards the erythroid and myeloid lineages (Fig. 3b). Differential expression analysis revealed that genes involved in the NF-κB signaling pathway, including *IL-1*β, as well as those involved in cardiovascular disorders, such as *CXCL2, CXCL3*, and *MIF*, were significantly downregulated in HSCs obtained at canakinumab response compared with those in HSCs obtained before treatment (Fig. 3c and Supplemental Fig. 3b,c), which aligns with previous findings that canakinumab can significantly reduce the risk of recurrent cardiovascular events in patients with previous myocardial infarction^30^. Given that HSCs expressed *IL-1*β and *IL-1R1* (Supplemental Fig. 3d), these data confirm on-target engagement of canakinumab. Moreover, consistent with previous findings showing that *DNMT3A* mutations provide HSCs with a competitive advantage under inflammatory conditions^31^, canakinumab decreased *DNMT3A* clonal burden at the time of erythroid hematological improvement after 2 cycles of canakinumab treatment (33.0% at baseline vs 24.9% at the end of cycle 2).

**Fig. 3.**
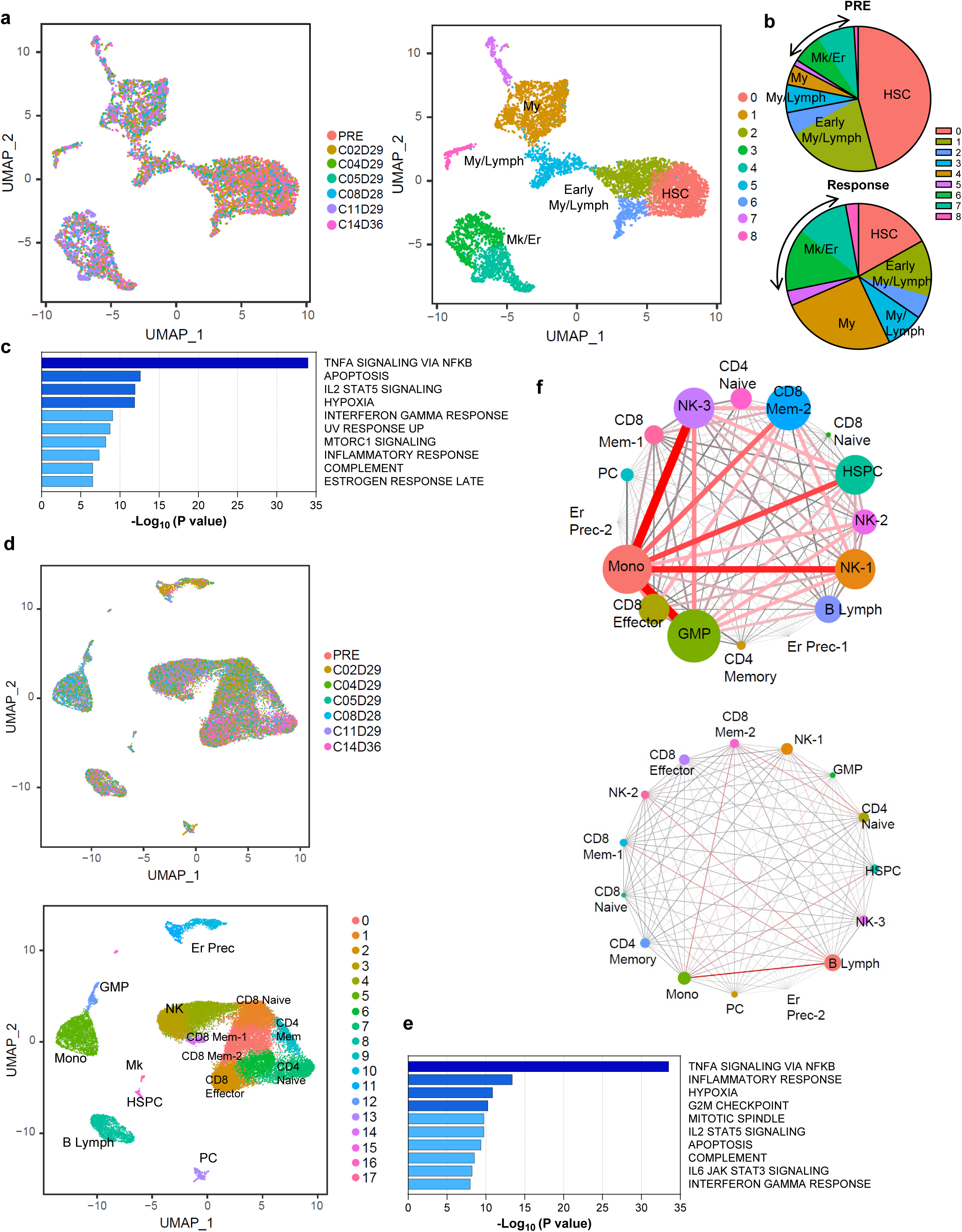
Targeting IL-1β pathway by canakinumab overcomes anemia in LR-MDS driven by *DNMT3A* and *TET2* mutations. **(a)** UMAP plot of scRNA-seq data from Lin^-^CD34^+^ cells isolated from the BM of the MDS patient UPN-02 before canakinumab treatment (PRE; n = 1,381) and after 2, 4, 5, 8, 11, and 14 cycles of canakinumab treatment (n = 5,721). Each dot represents one cell. Different colors represent the sample (left) and cluster (right) identities. My, myeloid; Lymph, lymphoid; HSC, hematopoietic stem cell; Mk, megakaryocytic; Er, erythroid. **(b)** Cluster distribution of Lin^-^ CD34^+^ cells isolated from the BM before canakinumab treatment (PRE) and during response to canakinumab treatment (cycles 2, 4, 5, 8, and 11), represented as the percentage of cells in each cluster shown in Fig. 3a. Clusters are grouped based on the cells’ lineage annotation. Black arrows indicate the Mk/Er clusters. **(c)** Pathway enrichment analysis of genes that were significantly (*P* ≤ 0.05) downregulated in HSCs after 2 cycles of canakinumab treatment compared with those in HSCs before canakinumab treatment. The top 10 Hallmark gene sets are shown. **(d)** UMAP plot of scRNA-seq data from BM MNCs isolated from patient UPN-02 before canakinumab treatment (PRE; n = 2,980) and after 2, 4, 5, 8, 11, and 14 cycles of canakinumab treatment (n = 27,913). Each dot represents one cell. Different colors represent the sample (top) and cluster (bottom) identities. Er, erythroid; Prec, precursor; GMP, granulo-monocytic progenitor; NK, natural killer; Mem, memory; Mono, monocytic; Mk, megakaryocytic; HSPC, hematopoietic stem and progenitor cell; Lymph, lymphoid; PC, plasma cell. **(e)** Pathway enrichment analysis of genes that were significantly (*P* ≤ 0.05) downregulated in monocytes after 2 cycles of canakinumab treatment compared with those in monocytes before canakinumab treatment. The top 10 Hallmark gene sets are shown. **(f)** Connectome web analysis of interacting cell types among MNCs isolated from the BM of patient UPN-02 before canakinumab treatment (top) and after 2 cycles of canakinumab treatment (bottom). The vertex (i.e., colored cell node) size is proportional to the number of interactions to and from each cell type, and the thickness of each connecting line is proportional to the number of interactions between 2 nodes. NK, natural killer; Mem, memory; PC, plasma cell; HSPC, hematopoietic stem and progenitor cell; Er, erythroid; Prec, precursor; Mono, monocytic; Lymph, lymphoid; GMP, granulo-monocytic progenitor.

ScRNA-seq analysis of MNCs obtained before and after canakinumab administration (Fig. 3d) revealed that the expression of major regulators of the NF-κB and inflammatory signaling pathways in MDS, including *TLR2, KDM6B*, *REL*, and *NLRP3,* were significantly downregulated in the monocyte population (Fig. 3e and Supplemental Fig. 3e), which also expressed *IL-1*β and *IL-1R1* (Supplemental Fig. 3f). Consistent with this observation, the levels of several pro-inflammatory cytokines, including IL-1β, IL-18, IL-6, and IFN-γ, were reduced in the BM plasma collected at the time of canakinumab response compared with those in the BM plasma collected before treatment (Supplemental Fig. 3g). This broad spectrum of pro-inflammatory cytokine inhibition was associated with the recovery of BM erythroblasts (Supplemental Fig. 3h) and significant changes in the immune microenvironment, including a decrease in the CD8^+^GZMK^+^ memory T cell population (Supplemental Fig. 3i-k), which is associated with several inflammatory conditions and further enhances the release of pro-inflammatory cytokines^32–34^. Consistent with this observation, when we inferred cell-to-cell communication from the combined expression of multi-subunit ligand–receptor complexes using CellPhoneDB, a repository of ligands and receptors and their interactions^35^, we observed that before canakinumab treatment, CD8^+^GZMK^+^ T cells were predicted to significantly interact with the monocyte population to drive these cells’ migration (e.g., a4b1:PLAUR; CCL3/CCL3L1/CCL5:CCR1) and differentiation into highly pro-inflammatory macrophages (e.g., IFN-γ:IFNR, LTB:LTBR; Fig. 3f and Supplemental Fig. 3l).

In contrast, canakinumab treatment did not rescue aberrant erythroid differentiation in LR-MDS patients with multiple genetic aberrations (Supplemental Table 11), as shown by scRNA-seq analyses of Lin^-^CD34^+^ HSPCs (Supplemental Fig. 3m) and MNCs (Supplemental Fig. 3n) isolated from 2 representative patients (UPN-7 and UPN-14) whose disease did not respond to canakinumab. Similar data were obtained for 2 other patients whose disease progressed after canakinumab therapy (data not shown). Given that *IL-1*β and *IL-1R1* were expressed in both HSCs and monocytes (Supplemental Fig. 3o,p) and that canakinumab significantly decreased NF-κB–mediated inflammatory signaling in these populations (Supplemental Fig. 3q,r), these data support our preclinical results that IL-1β-mediated inflammation is an initiating event in the development of clonal hematopoiesis, but it does not account for disease progression, when clonal complexity is acquired and other intrinsic (e.g., cooperative mutation effects) and extrinsic factors (e.g., exacerbation of immune suppression) contribute to clonal expansion of mutant cells.

In summary, we demonstrated that the most frequent CCUS mutations, *DNMT3A* and *TET2*, rewire the HSCs’ aging phenotype and remodel the BM microenvironment by upregulating IL-1β-driven inflammasome signaling, which leads to HSCs’ exit from quiescence and myeloid-skewing at the expense of erythropoiesis (Supplemental Fig. 3s). Consistent with these observations, targeting the IL-1β signaling pathway with canakinumab rescued anemia only in LR-MDS patients with single driver mutations in *DNMT3A* or *TET2*. These results align with unpublished data showing that canakinumab specifically improved hemoglobin levels in patients with clonal hematopoiesis driven by *DNMT3A* and *TET2* mutations^36^.

Importantly, canakinumab treatment also decreased the expression of genes involved in cardiovascular diseases, such as *CXCL2*, *CXCL*3, and *MIF* in LR-MDS with single mutations in *DNMT3A* or *TET2*, which confirmed previous studies showing increased efficacy of this agent in reducing recurrent cardiovascular events in patients with previous myocardial infarction carrying mutations in *TET2* compared with those without any mutation^37^.

Further clinical trials of canakinumab in patients with CCUS will clarify whether the modulation of IL-1β-induced-inflammation can improve these patients’ peripheral blood cytopenias and reduce their risk of developing cardiovascular disorders^38^, thus modifying the course of the disease.

## ACKNOWLEDGMENTS

This work was supported by philanthropic contributions to MD Anderson’s AML and MDS Moon Shot Program, by the P. Evans Foundation, and by a Women Who Conquer Cancer Young Investigator Award to K.C. This work used MD Anderson’s South Campus Flow Cytometry and Cell Sorting Core Laboratory, Orion Core, and Sequencing and Microarray Facility, all of which were supported in part by the National Institutes of Health (National Cancer Institute) through the MD Anderson’s Cancer Center Support Grant (P30 CA16672). S.C. is a Scholar of the Leukemia and Lymphoma Society. The authors thank Joseph Munch for assistance with manuscript editing.

## AUTHOR CONTRIBUTIONS

S.C. designed experiments. V.A., I.G.-G., H.Y., N.T., and R.T. contributed to the experiments; J.J.R.-S., K.C., G.M.-B., and D.H. analyzed the clinical data; R.K.-S. and S.L. analyzed the NGS data; F.M. and Y.G. analyzed the scRNA-seq data; L.T. and P.L. analyzed the metabolomic data; K.C.-D., G.A.-A., M.P., R.B., G.G.M., and D.H. made critical intellectual contributions throughout the project; G.G.M. ran the clinical trial; S.C. wrote the manuscript.

## DECLARATION OF INTERESTS

G.G.-M. reports clinical funding from Novartis, AbbVie, and Amgen. All other authors report no conflicts of interest relative to this work.

Correspondence and requests for materials should be addressed to.

## METHODS

### Human samples

BM specimens from 20 untreated patients with CCUS and 3 untreated patients with LR-MDS were obtained in accordance with the Declaration of Helsinki from the Department of Leukemia at MD Anderson Cancer Center (MDACC) under protocol PA15-0926 or from the Department of Medicine at University of California San Diego, with the approval of the corresponding Institutional Review Boards. BM samples from 3 yHDs (median age = 21 years, 33% male) and 26 eHDs (median age = 54 years, 58% male) were obtained from AllCells (Alameda, CA) or from the Department of Stem Cell Transplantation and Cellular Therapy at MDACC. Written informed consent was obtained from all donors, and all diagnoses were confirmed by dedicated hematopathologists. The clinical characteristics of the patients with available information are shown in Supplemental Table 1.

BM MNCs were isolated from each sample using a standard gradient separation approach with Ficoll-Paque PLUS (GE Healthcare Life Sciences, Pittsburgh, PA, USA). MNCs were enriched in CD34^+^ cells using magnetic separation with the CD34 Microbead Kit (Miltenyi Biotec, San Diego, CA, USA) and further purified by flow cytometric sorting as described below.

### Flow cytometry and fluorescence-activated cell sorting (FACS)

Quantitative flow cytometry and FACS of live human MNCs and CD34^+^ cells were performed using a previously described gating strategy and antigen panel^10^ and antibodies against CD2 (RPA-2.10; 1:20), CD3 (SK7; 1:10), CD14 (MφP9; 1:20), CD19 (SJ25C1; 1:10), CD20 (2H7; 1:10), CD34 (581; 1:20), CD56 (B159; 1:40), CD123 (9F5; 1:20), and CD235a (HIR2; 1:40; all from BD Biosciences); CD4 (S3.5; 1:20), CD11b (ICRF44; 1:20), CD33 (P67.6; 1:20), and CD90 (5E10; 1:10; all from ThermoFisher Scientific, Waltham, MA); CD7 (6B7; 1:20) and CD38 (HIT2; 1:20; both from BioLegend, San Diego, CA); CD10 (SJ5-1B4; 1:20; Leinco Technologies, St Louis, MO); and CD45RA (HI100; 1:10; Tonbo Biosciences, San Diego, CA).

Analyzed or FACS-purified samples were acquired with a BD Influx Cell Sorter (BD Biosciences), and the cell populations were analyzed using FlowJo v10.7.1 software (FlowJo, Ashland, OR). All experiments included fluorescence-minus-one and single-stained controls, and were performed at the Advanced Cytometry & Sorting Facility at MDACC.

### scRNA-seq

FACS-purified live BM MNCs or Lin^-^CD34^+^ cells were sequenced at the Advanced Technology Genomics Core facility at MDACC. Sample concentration and cell suspension viability were evaluated using a Countess II FL Automated Cell Counter (ThermoFisher Scientific) and manual counting. Samples were normalized for input into the Chromium Single Cell A Chip Kit (10x Genomics, Pleasanton, CA), in which single cells were lysed and barcoded for reverse-transcription. The pooled single-stranded, barcoded cDNA was amplified and fragmented for library preparation. Pooled libraries were sequenced using a NovaSeq6000 SP 100-cycle flow cell (Illumina, San Diego, CA).

Sequencing analysis was performed using 10X Genomics’ CellRanger software, version 3.0.2. Fastq files were generated using CellRanger MkFastq pipeline (version 3.0.2). Raw reads were mapped to the human reference genome (refdata-cellranger-GRCh38-3.0.0) using the CellRanger Count pipeline. Multiple samples were aggregated using the Cellranger Aggr pipeline. The digital expression matrix was analyzed using the R package Seurat (version 3.0.2)^39^ to identify different cell types and signature genes for each. Cells with fewer than 500 unique molecular identifiers or greater than 50% mitochondrial expression were excluded from further analysis. The Seurat function NormalizeData was used to normalize the raw counts. Variable genes were identified using the FindVariableFeatures function. The ScaleData function was used to scale and center expression values in the dataset, and the number of unique molecular identifiers was regressed against each gene. Uniform manifold approximation and projection (UMAP) was used to reduce the dimensions of the data, and the first two dimensions were used in the plots. FindClusters function was used to cluster the cells. Marker genes for each cluster were identified using the FindAllMarker function. Cell types were annotated based on marker genes and canonical markers^4–7^. Pathway analyses of differentially expressed genes were conducted using Metascape^40^.

CellphoneDB (v2.0.0)^35^ was used to analyze ligand–receptor interactions. Briefly, each cell type was separated by disease classifications, and a separate run was performed for each disease classification. The connectome web was plotted using the igraph package in R.

### Metabolomic analysis using ultra-high resolution mass spectrometry

CD34^+^ cells were isolated by magnetic enrichment as indicated above, washed with ice-cold PBS, and pelleted by centrifugation at 1,000 × *g* for 2 min at 4°C. Cell pellets were snap-frozen in liquid nitrogen and temporarily stored at −80°C. Sample analysis was carried out in the Metabolomics Facility at MDACC. Metabolites were extracted from cell pellets by adding 1 mL ice-cold 0.1% ammonium hydroxide in 80/20 (v/v) methanol/water. Extracts were centrifuged at 17,000 × *g* for 5 min at 4°C, and the supernatants were transferred to clean tubes and evaporated to dryness under nitrogen. Dried extracts were reconstituted in deionized water, and 10 μL was injected for analysis by ion chromatography–mass spectrometry. Ion chromatography mobile phase A (weak) was water, and mobile phase B (MPB; strong) was water containing 100 mM KOH. Ion chromatography was performed using a Dionex ICS-5000^+^ system (ThermoFisher Scientific), which included a Thermo IonPac AS11 column (4-µm particle size, 250 × 2 mm) with the column compartment kept at 30°C. The autosampler tray was cooled to 4°C. The mobile phase flow rate was 360 µL/min, and the gradient elution program was 0-5 min, 1% MPB; 5-25 min, 1-35% MPB; 25-39 min, 35-99% MPB; 39-49 min, 99% MPB; 49-50 min, 99-1% MPB. The total run time was 50 min. For better sensitivity, desolvation was assisted by methanol, which was delivered by an external pump and combined with the eluent via a low dead volume mixing tee. Data were acquired using a Thermo Orbitrap Fusion Tribrid Mass Spectrometer under ESI negative ionization mode at a resolution of 240,000. Raw data files were imported into Thermo Trace Finder software for the final analysis. The relative abundance of each metabolite was normalized to the numbers of live cells. Levels of 1,3-BPG and 3-PG were inferred from the amount of their isomers 2-phospho-glycerate and 2,3-diphosphoglycerate, respectively.

### Quantification of cytokines, chemokines, and growth factors in BM plasma

EGF, eotaxin, G-CSF, GM-CSF, IFN-α2, IFN-γ, IL-1α, IL-1β, IL-1RA, IL-2, IL-3, IL-4, IL-5, IL-6, IL-7, IL-8, IL-10, IL-12 (p40), IL-12 (p70), IL-13, IL-15, IL-17A, IL-17E/IL-25, IL-17F, IL-18, IL-22, IP-10, MCP-1, M-CSF, MIG, MIP-1α, MIP-1β, PDGF-AA, PDGF-AB/BB, TNF-α, TNF-β, VEGF-A, and RANTES were simultaneously quantified from the plasma collected from the BM of patients enrolled in the clinical trials using the MILLIPLEX Human Cytokine/Chemokine/Growth Factor Panel A (HCYTA-60K-PX38, Millipore) according to the manufacturer’s guidelines. Samples were assayed neat, and 50 µL of a 1:1 mixture of the sample and assay buffer was added to each assay plate well. For the detection of RANTES, the samples were initially diluted with the assay buffer to 1:100. Briefly, 25 µL of the 38-analyte bead mixture was added to each plate well, and the plate was incubated overnight at 4°C. The following day, the plate was incubated at room temperature for 30 min with shaking at 500 rpm. After three washes, 25 µL of detection antibody was added to each well, and the plate was incubated for 1 h at room temperature with shaking at 500 rpm. Then, 25 µL of streptavidin–phycoerythrin solution was added to each well, and the plate was incubated for 30 min at room temperature with shaking at 500 rpm. After three wash steps, 150 µL of xMAP Sheath Fluid Plus (4050021, ThermoFisher) was added to each well. Acquisition was performed using the Luminex 200 system and the xPONENT version 4.2 software. Analysis was performed using the Bio-Plex Manager version 6.1 software.

### Clinical trial

Clinical data were collected retrospectively by reviewing the patients’ electronic health records. This study was conducted at MDACC and approved by the Institutional Review Board in accordance with the Declaration of Helsinki. Written informed consent was obtained from all the patients. The study was registered at ClinicalTrials.gov (NCT04239157). Canakinumab was supplied by Novartis (Basel, Switzerland) and was administered subcutaneously once every 28 days. BM aspiration and/or biopsy, along with flow cytometric, cytogenetic, and mutation analyses using an 81-gene next-generation sequencing panel^41^, were performed before therapy, on day 28 (± 3 days) of cycle 2, and then every 3 cycles thereafter or more frequently as clinically warranted. Canakinumab was discontinued after 6 cycles if it failed to elicit a response but was otherwise continued until disease progression. Responses were assessed using the modified International Working Group 2006 criteria for MDS^42^. Overall response included complete response, marrow complete response, hematological improvement, and a combination of marrow complete response and hematological improvement. The clinical characteristics of patients enrolled in the canakinumab trial are shown in Supplemental Table 11.

### Experimental data analysis

Quantitative data were analyzed using the GraphPad Prism 10 software (GraphPad, La Jolla, CA). Figure legends indicate the statistical tests used in each experiment. Statistically significant differences in the figures are indicated as \**P* < 0.05, \*\**P* < 0.01, \*\*\**P* < 0.001, and \*\*\*\**P* < 0.0001. Supplemental Fig. 3a,s were made using Biorender.com.

## Data availability

Data sets generated in this study by scRNA-seq are accessible at GEO (GSE237148).

---

**Supplemental Fig. 1. *DNMT3A* and *TET2* mutations transcriptionally rewire the HSCs’ aging phenotype.**

**(a)** Expression of a curated set of lineage marker genes in the 16 scRNA-seq clusters shown in Fig. 1a. **(b)** Expression of *MLLT3*, *HLF*, *MEG3,* and *CDKN1C* in Lin^-^CD34^+^ cells. **(c)** Left, frequencies of immunophenotypically-defined HSPC populations in Lin^-^CD34^+^ cells from BM samples of the same yHDs (n = 3) and eHDs (n = 4) analyzed by scRNA-seq (Fig. 1a). Each symbol represents a single sample. Bars represent means ± standard deviations (SDs). Statistically significant differences were determined using the Student’s t-test (\**P* ≤ 0.05; \*\**P* ≤ 0.01). Right, flow cytometry plots of gating strategy for CD34^+^-enriched BM cells from representative yHDs and eHDs. Percentages indicate cell frequencies in the corresponding gates. LT-HSC, long-term HSC; MPP, multipotent progenitor; LMPP, lymphoid-primed MPP; CMP, common myeloid progenitor; GMP, granulo-monocytic progenitor; MEP, megakaryocytic-erythroid progenitor. **(d)** Dot plot of the expression levels of genes involved in TNF-α signaling through NF-κB that were significantly (*P* adj ≤ 0.05) upregulated in HSCs from eHDs compared with those in HSCs from yHDs. **(e)** Dot plot of the expression levels of genes involved in IFN-γ signaling that were significantly (*P* adj ≤ 0.05) upregulated in HSCs from eHDs compared with those in HSCs from yHDs. **(f)** *MCL1* expression in HSCs from yHDs and eHDs. Each dot represents the expression level of a single cell (*P* adj = 3.71 × 10^-14^). **(g)** UMAP plots of scRNA-seq data from Lin^-^CD34^+^CD38^-^CD90^+^CD45RA^-^ LT-HSCs isolated from 3 yHDs (n = 3,215) and 2 eHDs (n = 1,759). Each dot represents one cell. Different colors represent the sample (left) and cluster (right) identities. **(h)** Pathway enrichment analysis of genes that were significantly (*P* adj ≤ 0.05) upregulated in LT-HSCs from eHDs compared with those in LT-HSCs from yHDs. The top 10 Hallmark gene sets are shown. **(i)** Dot plot of the expression levels of genes involved in TNF-α signaling through NF-κB which were significantly (*P* adj ≤ 0.05) upregulated in LT-HSCs from eHDs compared with those in LT-HSCs from yHDs. **(j)** Dot plot of the expression levels of genes involved in IFN-γ signaling that were significantly (*P* adj ≤ 0.05) upregulated in LT-HSCs from eHDs compared with those in LT-HSCs from yHDs. **(k)** Frequency of Lin^-^ CD34^+^ CD38^-^ HSCs (left) and Lin^-^CD34^+^CD38^+^ myeloid progenitor cells (right) in total BM MNCs from 18 eHDs and 14 CCUS patients. Each symbol represents a single sample. The lines represent the median ± interquartile range. \*\*\*\**P* < 0.0001, Mann-Whitney test. **(l)** Dot plot of the expression levels of genes involved in translation that were significantly (*P* adj ≤ 0.05) upregulated in HSCs from CCUS patients compared with those in HSCs from eHDs. **(m)** Dot plot of the expression levels of genes involved in respiratory electron transport that were significantly (*P* adj ≤ 0.05) upregulated in HSCs from CCUS patients compared with those in HSCs from eHDs. **(n)** Dot plot of the expression levels of genes involved in mitochondrial translation initiation that were significantly (*P* adj ≤ 0.05) upregulated in HSCs from CCUS patients compared with those in HSCs from eHDs. **(o)** Differences in the amounts of soluble metabolites from glycolysis and tricarboxylic acid (TCA) cycle between CD34^+^ HSPCs from CCUS patients (n = 5) and CD34^+^ HSPCs from eHDs (n = 6). Quantitative differences are shown as the log_10_ of the fold-change in the metabolite amount in HSPCs from CCUS patients compared with that in HSPCs from eHDs (left) and are represented in a metabolic pathway scheme (right). Light typography and dashed lines indicate decreased amounts of a given metabolite, and bold typography indicates increased amounts. Metabolites with a white background were either below the detection range in HDs or were not detected owing to method limitations. \**P* ≤ 0.05, \*\**P* ≤ 0.01, Student t-test. **(p)** Expression levels of *IL-1R1* in Lin^-^CD34^+^ cells shown in Fig. 1a. **(q)** Expression of *CXCL2*, *CXCL3,* and *MIF* in HSCs from eHDs and CCUS patients. Each dot represents the expression level of a single cell (*P* adj = 0.00047, *P* adj = 1.31 × 10^-11^, and *P* adj = 1.1 × 10^-4^, respectively).

**Supplemental Fig. 2. IL-1β–mediated inflammatory signaling is an initiation event in the pathogenesis of CCUS.**

**(a)** Expression of cluster marker genes among the 26 scRNA-seq clusters shown in Fig. 2a. The top 3 genes for each cluster are shown. **(b)** Lineage distribution of BM MNCs isolated from yHDs, eHDs and CCUS patients shown in Fig. 2a, represented as the percentage of cells in each cluster. HSPC, hematopoietic stem and progenitor cell; Myelo, myelomonocytic; Lymph, lymphoid; NK, natural killer; Er, erythroid; Prec, precursor; DC, dendritic cell. **(c)** Pathway enrichment analysis of genes significantly (*P* adj ≤ 0.05) upregulated in myelomonocytic cells from eHDs compared with myelomonocytic cells from yHDs. The top 10 Hallmark gene sets are shown. **(d)** Dot plot of the expression levels of genes involved in TNF-α signaling via NF-κB (left) and inflammatory response (right) which were significantly (*P* adj ≤ 0.05) differentially expressed between the myelomonocytic cells (Fig. 2a) obtained from yHDs and eHDs. **(e)** Expression of *CASP1* in the myelomonocytic cells from eHDs and CCUS patients shown in Fig. 2a (*P* adj = 7.30 × 10^-42^). **(f)** Expression of *S100A* genes in the myelomonocytes from eHDs and CCUS patients shown in Fig. 2a (*S100A4*, *P* adj = 5.05 × 10^-131^; *S100A6*, *P* adj = 6.83 × 10^-134^; *S100A8, P* adj = 4.62 × 10^-77^*; S100A9*, *P* adj = 4.40 × 10^-130^; *S100A10, P* adj = 1.57 × 10^-17^*; S100A11*, *P* adj = 1.82 × 10^-69^; *S100A12*, *P* adj = 2.29 × 10^-82^). **(g)** Expression of *MIF* (*P* adj = 2.14 × 10^-98^), *CXCL8* (*P* adj = 2.68 × 10^-20^), and *CXCL2* (*P* adj = 1.15 × 10^-20^) in HSCs from patients with CCUS and HSCs from patients with LR-MDS. Each dot represents the expression level of a single cell. **(h)** UMAP plot of scRNA-seq data from BM MNCs isolated from 3 patients with CCUS patients (n = 13,644) and 3 patients with LR-MDS (n = 12,346). Each dot represents a single cell. Different colors represent the sample (left) and cluster (right) identities. Er, erythroid; Prec, precursor; Mem, memory; HSPC, hematopoietic stem and progenitor cell; My, myeloid; Prog, progenitor; Mono, monocytic; PC, plasma cell; NK, natural killer; Lymph, lymphoid. **(i)** Expression of *IL-1*β (*P* adj = 2.69 × 10^-13^; left) and *CASP1* (*P* adj = 1.75 × 10^-7^; right) in monocytes from CCUS patients and LR-MDS patients. Each dot represents the expression level in a single cell.

**Supplemental Fig. 3. Targeting IL-1β pathway by canakinumab overcomes anemia in LR-MDS driven by *DNMT3A* and *TET2* mutations.**

(**a**) Schematic of UPN-01 and UPN-02 hemoglobin and platelet levels, and number of packed red blood cell (PRBC) units received before and after enrollment in the canakinumab trial. The black arrow indicates the response duration to canakinumab. C, cycle. **(b)** Dot plot of the expression levels of genes involved in TNF-α signaling through NF-κB which were significantly (*P* ≤ 0.05) differentially expressed between the HSCs (Fig. 3a) obtained before and after canakinumab treatment. **(c)** *MIF* expression in HSCs from patient UPN-02 before canakinumab treatment (PRE) and at the end of cycle 2 (C2) of canakinumab treatment. Each dot represents the expression level in a single cell (*P* adj = 1 × 10^-4^). **(d)** Expression levels of *IL-1β* (top) and *IL-1R1* (bottom) in Lin^-^CD34^+^ cells from patient UPN-02. **(e)** Dot plot of the expression levels of genes involved in TNF-α signaling through NF-κB (left) and inflammatory signaling (right) pathways that were significantly (*P* ≤ 0.05) differentially expressed between monocytes (Fig. 3d) obtained before and after canakinumab treatment. **(f)** Expression levels of *IL-1β* (top) and *IL-1R1* (bottom) in BM MNCs from patient UPN-02 before canakinumab treatment. **(g)** BM plasma concentrations of cytokines (pg/mL) that significantly decreased after cycle 2 (C2) of canakinumab treatment in patient UPN-02. Replicate raw values are plotted. Lines represent the median ± interquartile range. \**P* ≤ 0.05, \*\**P* ≤ 0.01, \*\*\**P* ≤ 0.001, \*\*\*\**P* < 0.0001, two-tailed Student t-test with Welch’s correction. **(h)** Cluster distribution of BM MNCs before canakinumab treatment (PRE) and at the end of cycle 2 (C2) and cycle 14 (C14) of canakinumab treatment, represented as the percentage of cells in each cluster. Arrows indicate the erythroblast populations. Clusters were grouped based on the cell lineage annotation. HSPC, hematopoietic stem and progenitor cell; Mk, megakaryocytic; GMP, granulocyte-monocyte progenitor; Mono, monocytic; Er, erythroid; Prec, precursor; NK, natural killer; Lymph, lymphoid; PC, plasma cell. **(i)** Expression levels of *CD8A* in MNCs isolated from the BM of patient UPN-02 before canakinumab treatment (PRE; left) and at the end of cycle 2 (C2; middle) and cycle 14 (C14; right) of canakinumab treatment. **(j)** Expression levels of *GZMK* in MNCs isolated from patient UPN-02 before canakinumab treatment (PRE; left) and at the end of cycle 2 (C2; middle) and cycle 14 (C14; right) of canakinumab treatment. **(k)** Cluster distribution of T-cell subtypes in the BM T-cell populations before canakinumab treatment (PRE; top) and at the end of cycle 2 (C2; middle) and cycle 14 (C14; right) of canakinumab treatment, represented as the percentage of cells in each cluster. Black arrows indicate T cells that represent the CD8^+^GZMK^+^ T-cell population. Clusters were grouped based on the cell lineage annotation. **(l)** Circle plot showing the most significant ligand-to-receptor interactions between CD8^+^GZMK^+^ T cells and monocytes that were gained from before canakinumab treatment to the end of cycle 2 (C2) of canakinumab treatment. **(m)** UMAP plot of scRNA-seq data from Lin^-^CD34^+^ cells isolated from the BM of MDS UPN-07 and UPN-14 patients before canakinumab treatment (PRE; n = 3,814) and after 2 cycles (C2) of canakinumab treatment (n = 4,043). Each dot represents a single cell. Different colors represent the sample (left) and cluster (right) identities. Lymph, lymphoid; My, myeloid; Er, erythroid; HSC, hematopoietic stem cell; Mk, megakaryocytic. **(n)** UMAP plot of scRNA-seq data from BM MNCs isolated from UPN-07 and UPN-14 patients before canakinumab treatment (PRE; n = 6,959) and after 2 cycles (C2) of canakinumab treatment (n = 6,977). Each dot represents one cell. Different colors represent the sample (top) and cluster (bottom) identities. Er, erythroid; Prec, precursor; NK, natural killer; Mk, megakaryocytic; Mem, memory; HSPC, hematopoietic stem and progenitor cell; Mono, monocytic; Lymph, lymphoid. **(o)** Expression levels of *IL-1β* (left) and *IL-1R1* (right) in Lin^-^CD34^+^ cells from patients UPN-07 and UPN-14. (**p)** Expression levels of *IL-1β* (left) and *IL-1R1* (right) in BM MNCs from patients UPN-07 and UPN-14. **(q)** Pathway enrichment analysis of genes significantly (*P* ≤ 0.05) downregulated in HSCs from patients UPN-7 and UPN-14 at the end of cycle 2 of canakinumab treatment compared with those in HSCs before canakinumab treatment. The top 10 Hallmark gene sets are shown. **(r)** Pathway enrichment analysis of genes significantly (*P* ≤ 0.05) downregulated in monocytes from patients UPN-7 and UPN-14 at the end of cycle 2 of canakinumab treatment compared with those in monocytes before canakinumab treatment. The top 10 Hallmark gene sets are shown. **(s)** Proposed working model. Physiological HSC aging is characterized by a progressive megakaryocytic bias. Aging-related somatic mutations, such as *DNMT3A* and *TET2*, rewire the HSCs’ aging phenotype. Mutant HSCs remodel the microenvironment by upregulating inflammatory genes mainly driven by IL-1β, which induces HSCs’ exit of quiescence and myeloid-skewed differentiation. Clonal myeloid-biased HSCs harboring *DNMT3A* and *TET2* mutations expand in the BM of CCUS patients and drive peripheral blood cytopenias. Inhibiting IL-1β–mediated inflammatory signaling with canakinumab can restore normal hematopoiesis and overcome transfusion dependency in patients with CCUS or early-stage MDS with *DNMT3A* or *TET2* mutations.

