## Supplementary figures and images for "IL-1β–mediated inflammatory signaling drives ineffective erythropoiesis in early-stage myelodysplastic syndromes"

### Supplementary Figures

#### Suppl Figure 1

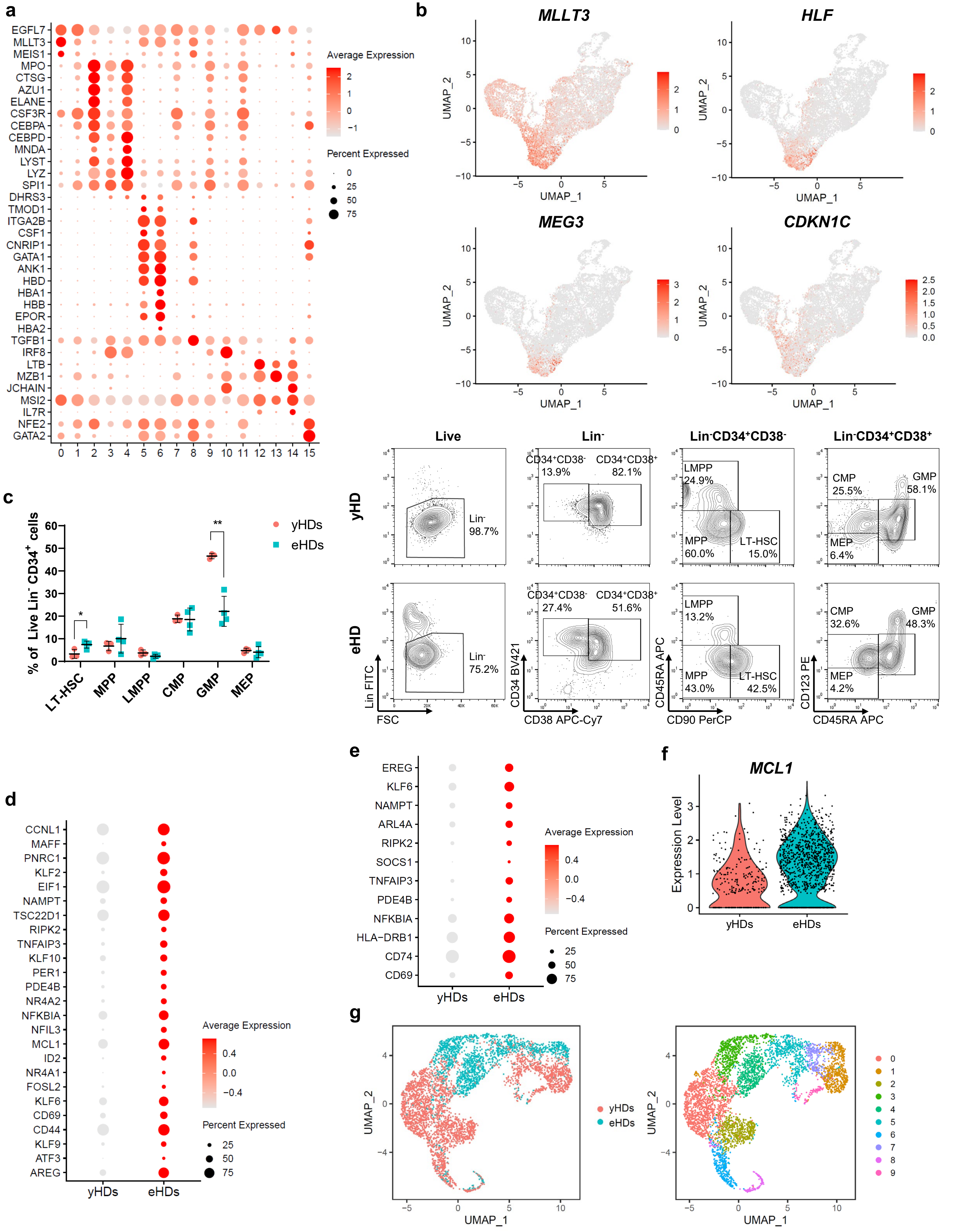

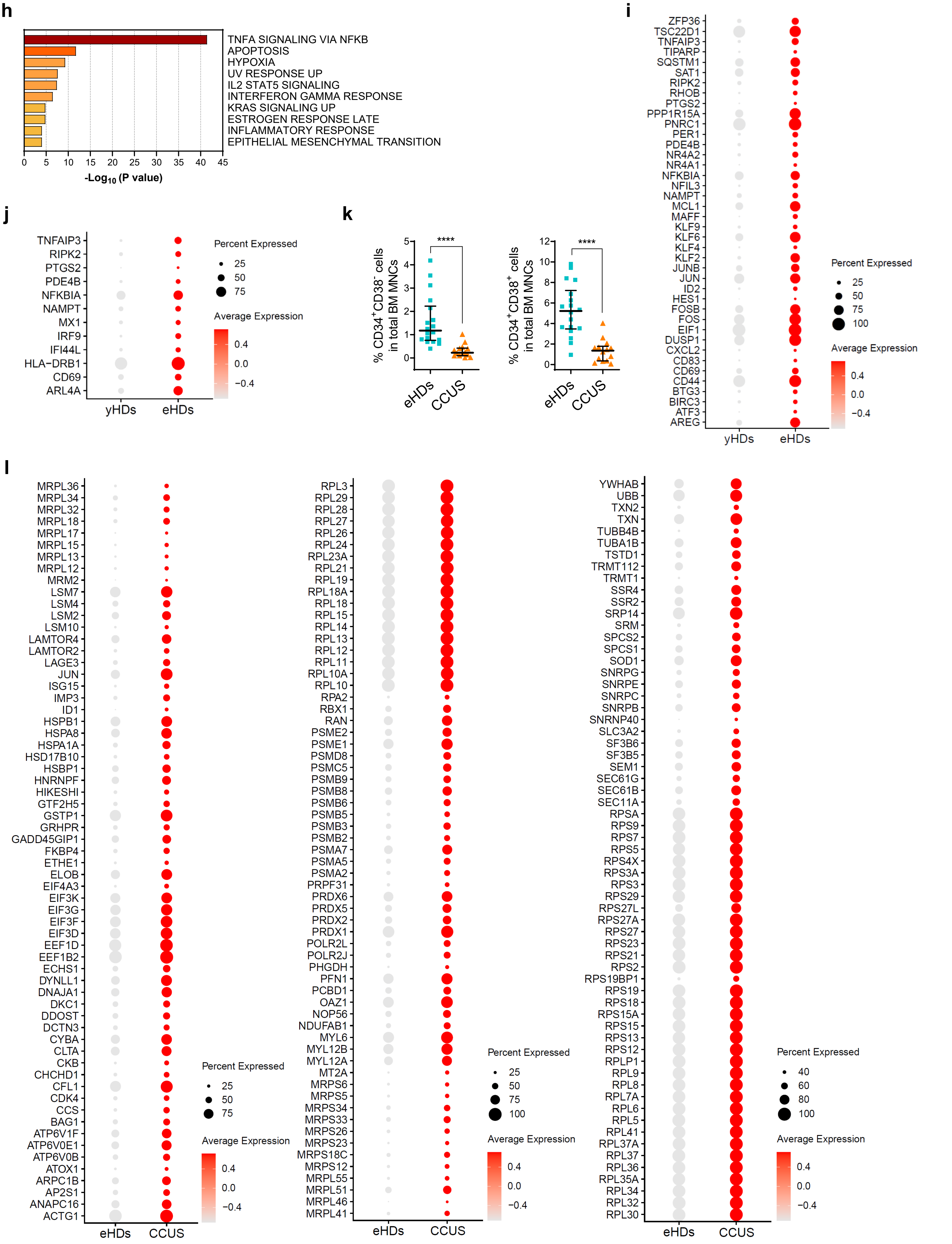

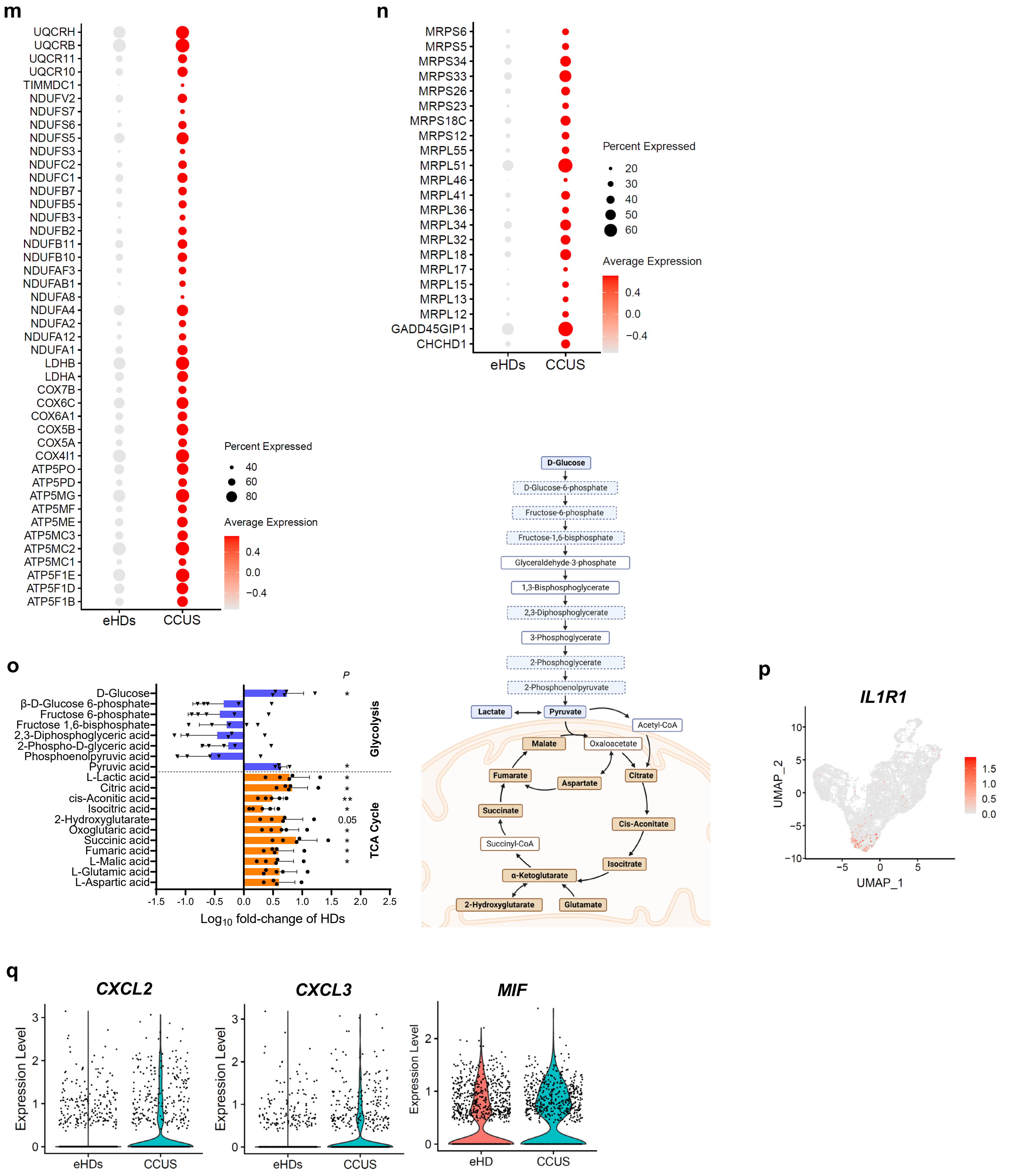

#### Suppl Figure 2

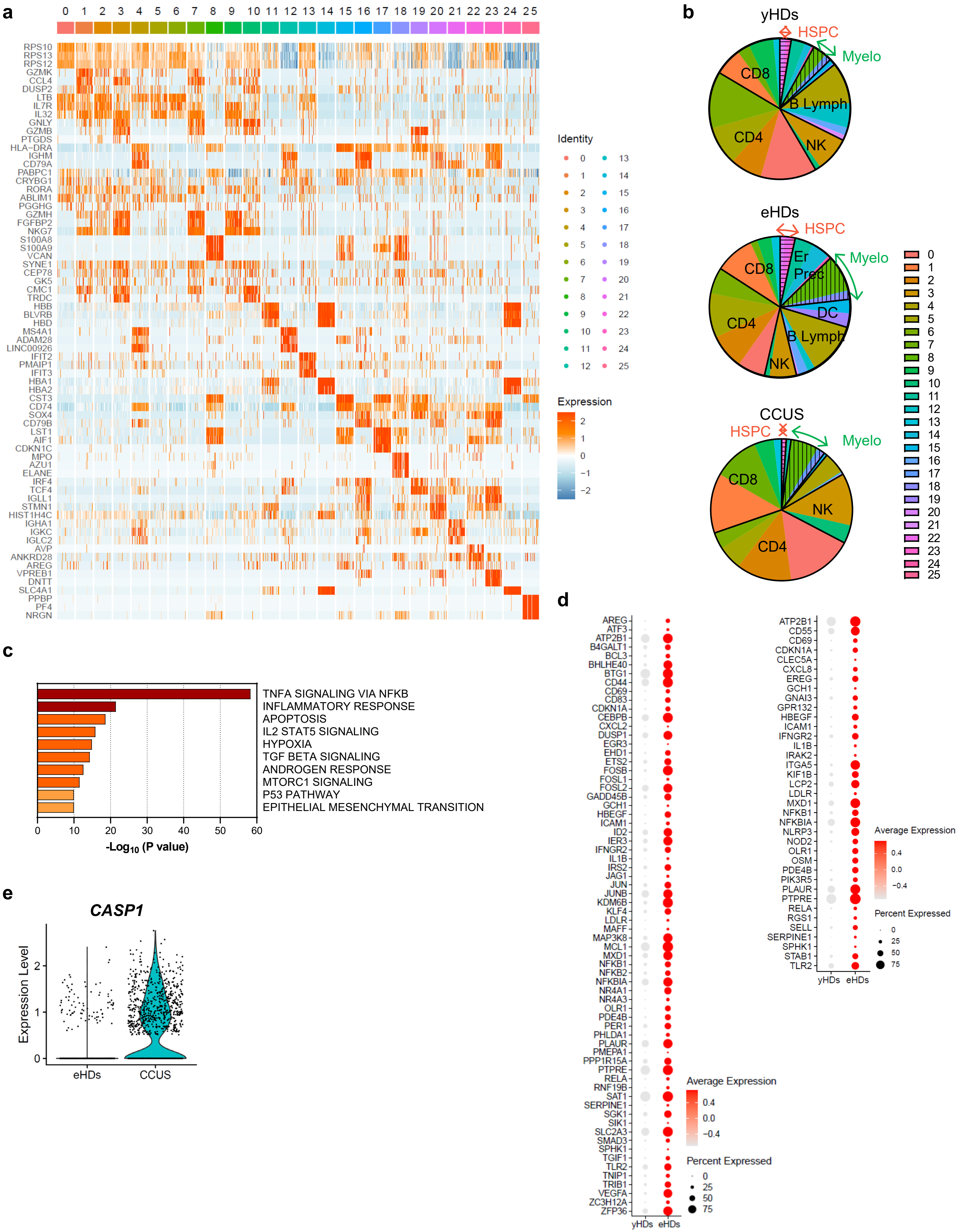

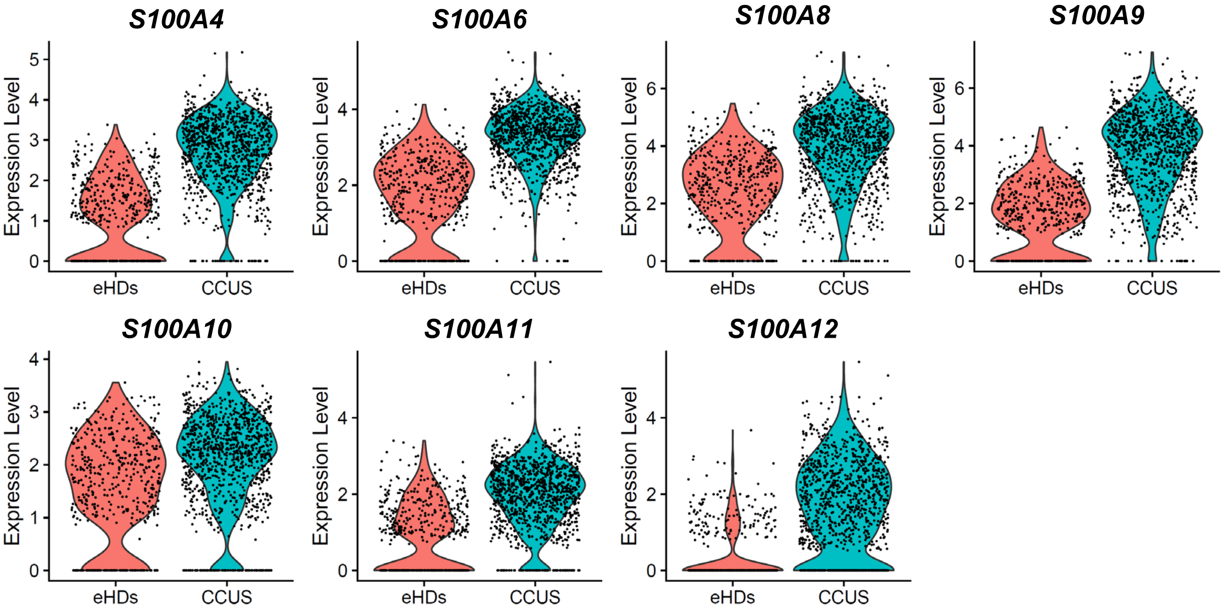

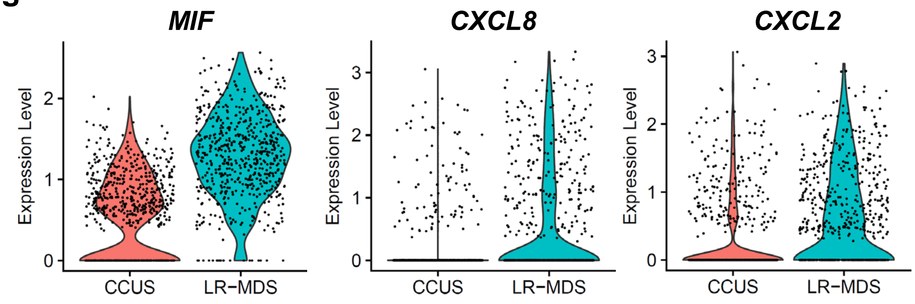

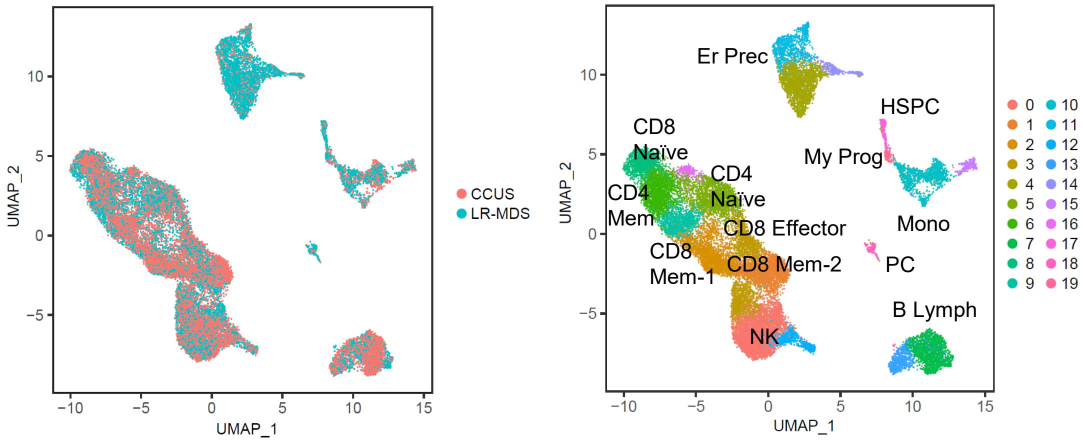

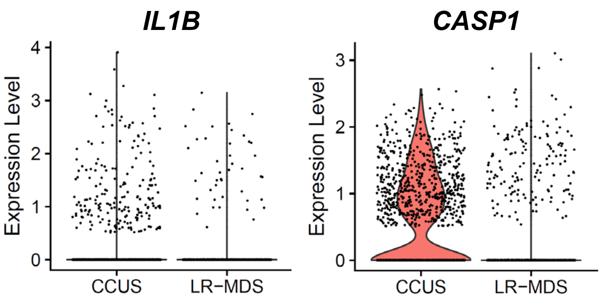

#### Suppl Figure 3

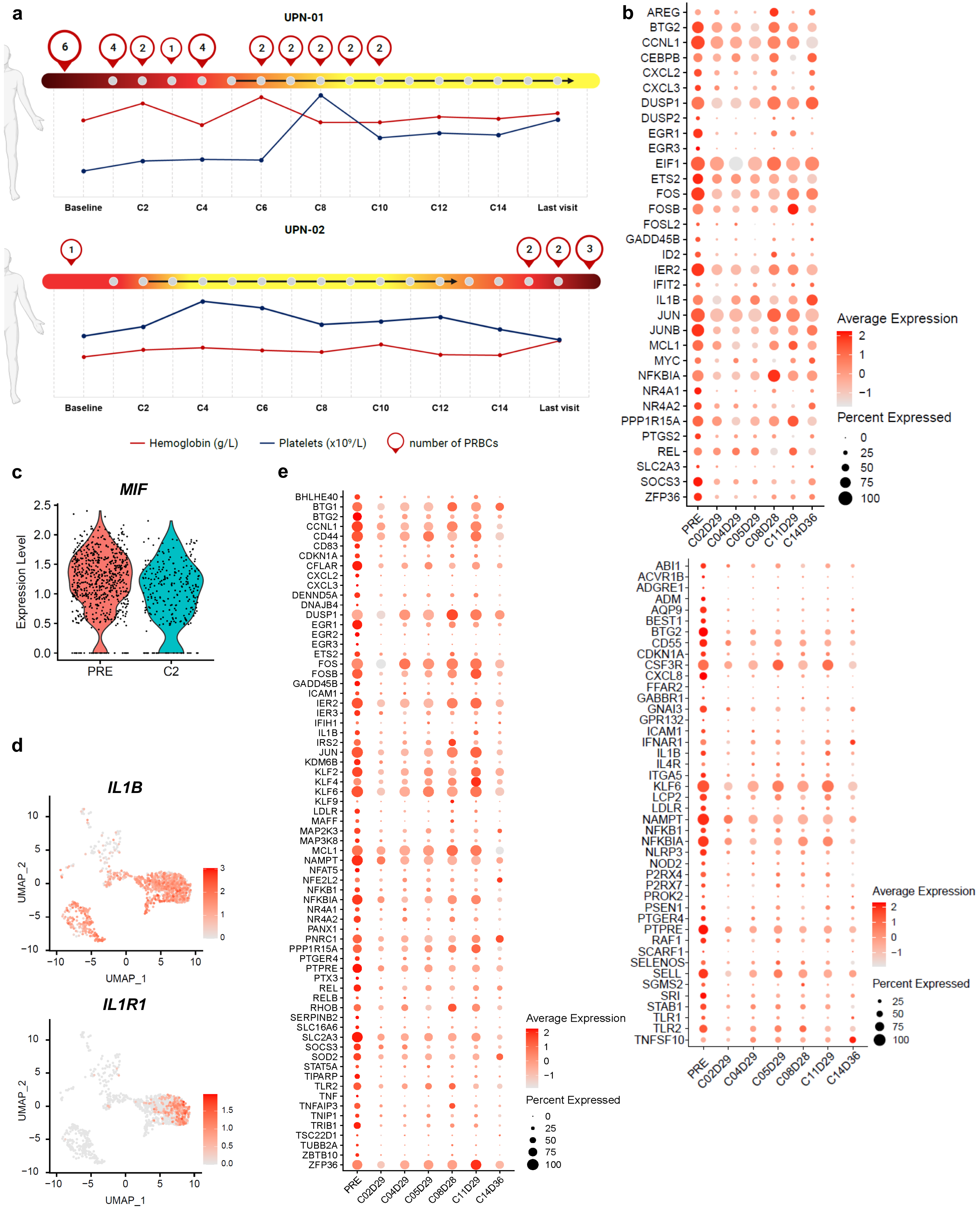

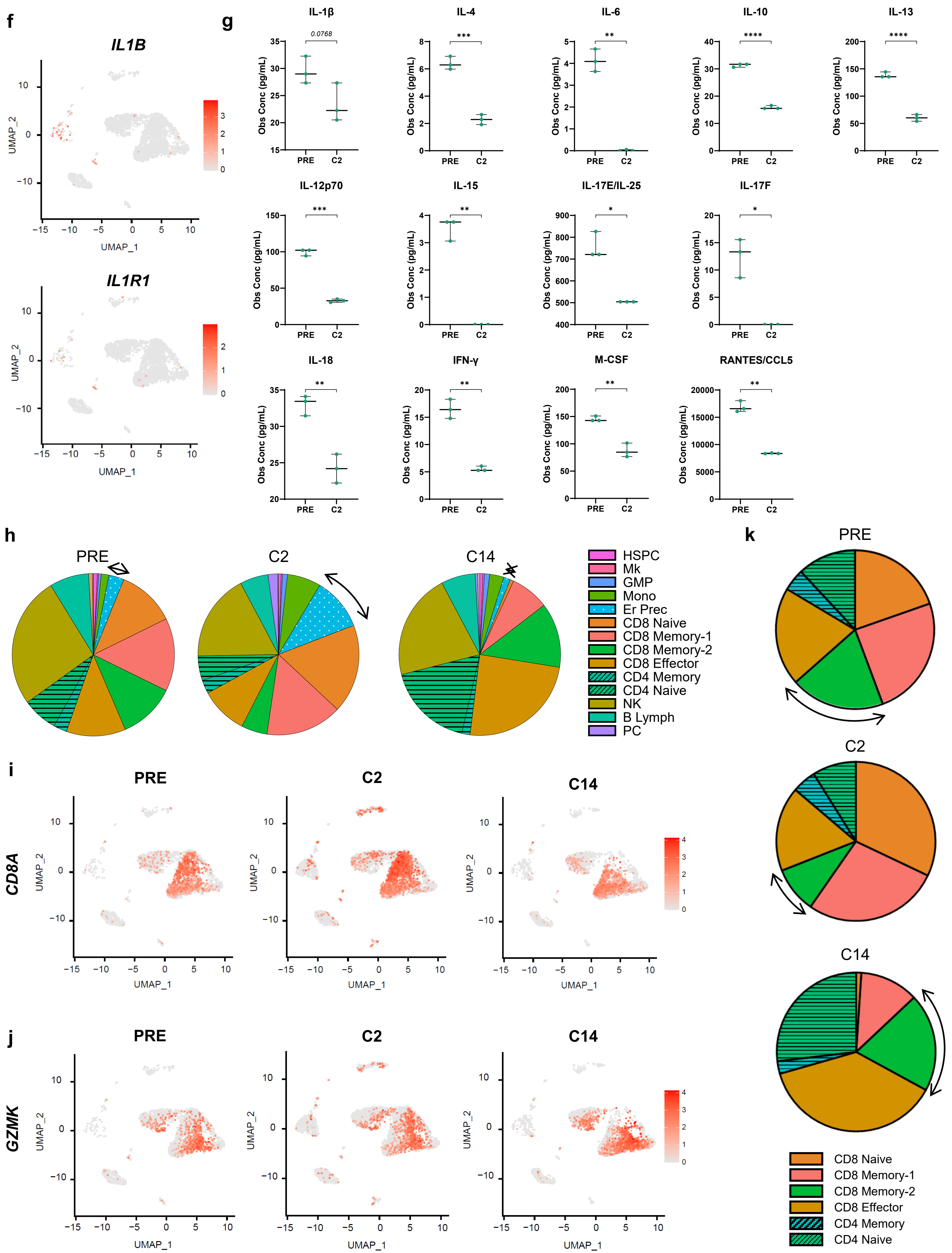

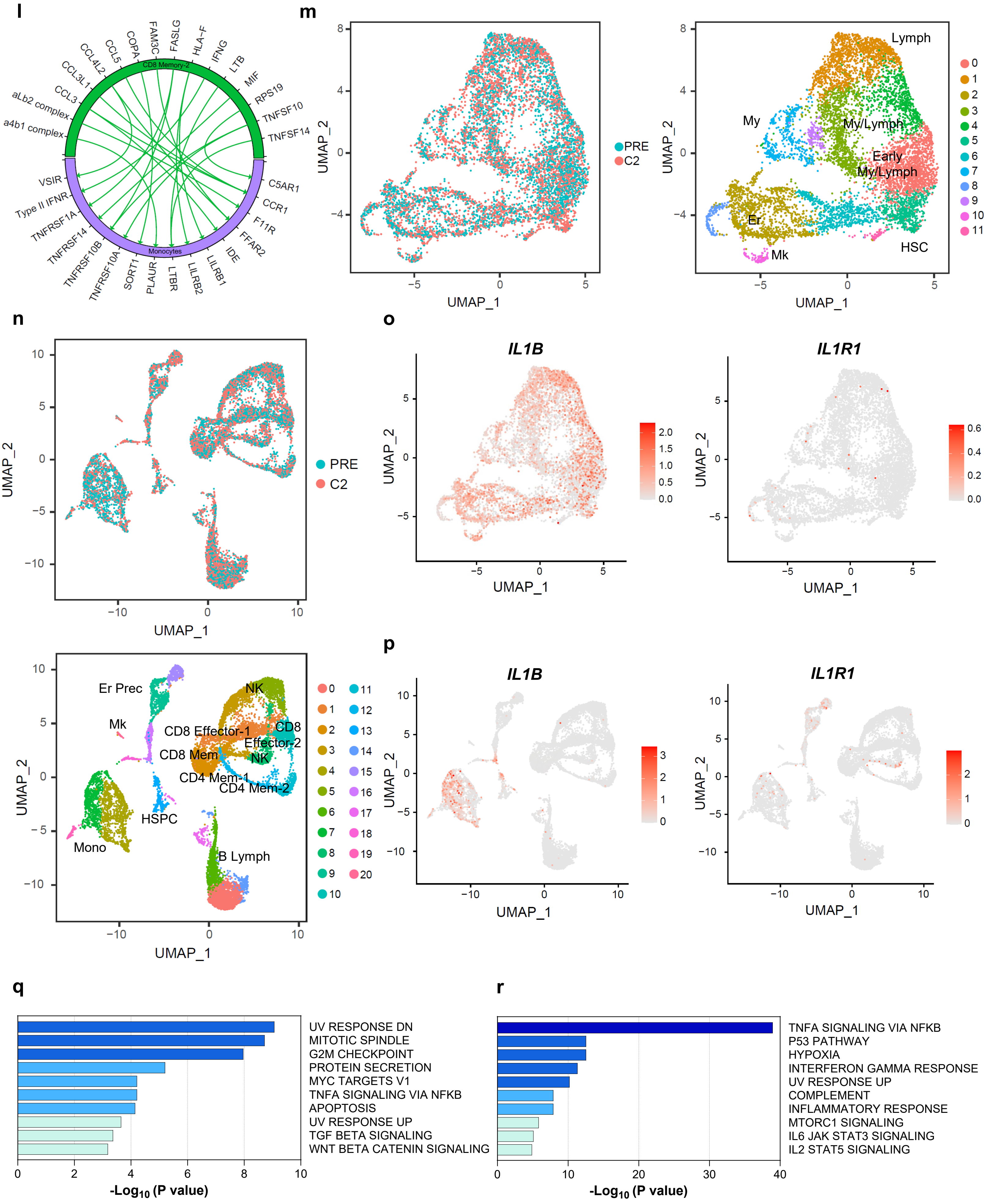

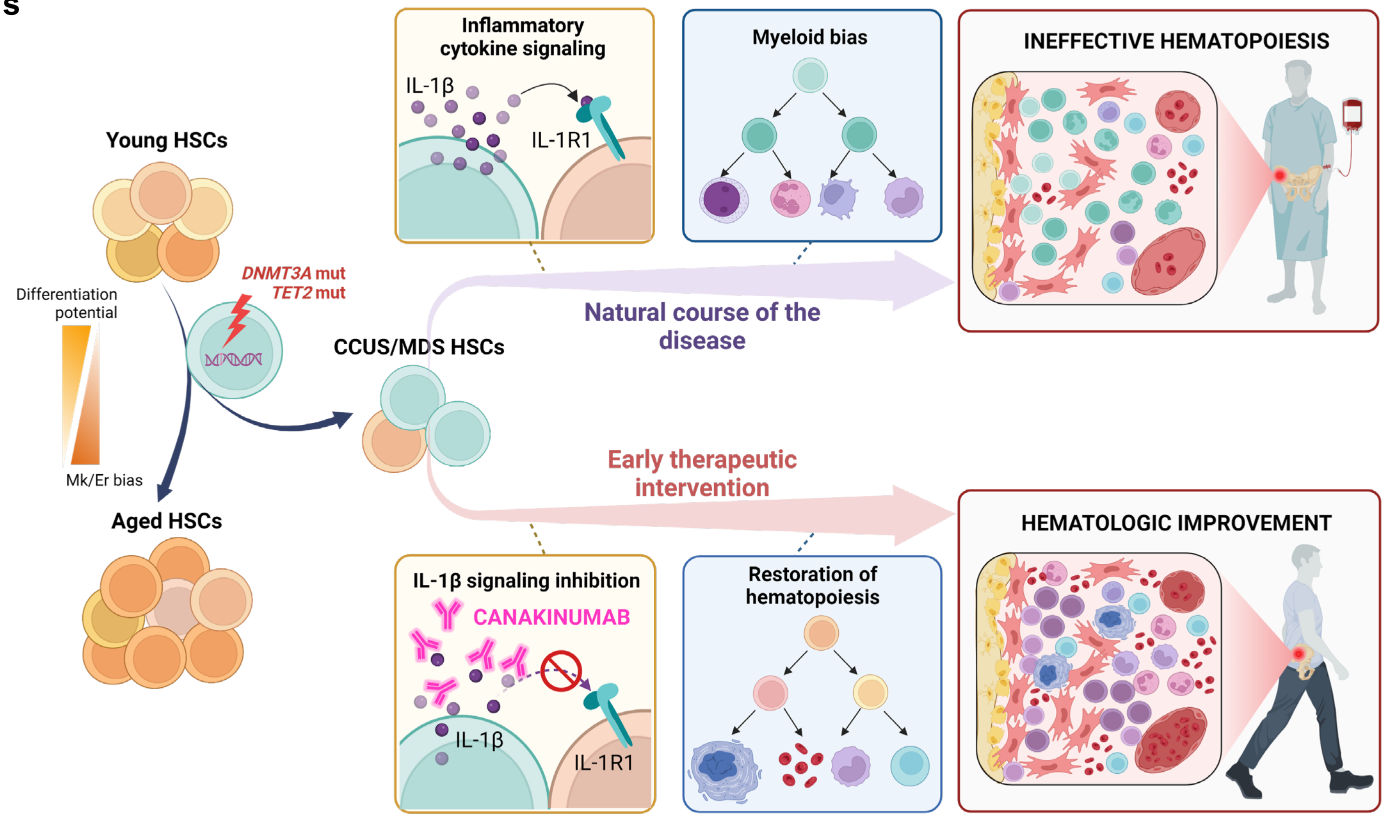
